# Ecological and Stochastic Determinants of the Growth and Persistence of the Oral Pathogen Porphyromonas gingivalis

**DOI:** 10.1101/2025.11.08.687400

**Authors:** Moemen Hussein, Arnab Barua, Mohammad Qasaimeh, Matthew Smardz, Patricia I. Diaz, Haralampos Hatzikirou

## Abstract

Population density plays a critical role in microbial fitness, yet its influence on pathogen colonization and persistence remains incompletely understood. *Porphyromonas gingivalis* (Pg) exhibits Allee-type growth, requiring a quorum threshold to replicate, yet is frequently detected at low abundance in vivo. We integrate quantitative growth experiments with mathematical modeling to identify ecological and stochastic determinants of Pg persistence. A cubic Allee-effect model quantifies a quorum threshold below which populations collapse, while conditioned medium from *Veillonella parvula* (Vp) lowers this threshold, indicating early-colonizer facilitation. Stochastic extensions and Fokker–Planck analysis show that microenvironmental noise enables escapes across the Allee barrier, consistent with long-term subthreshold experiments yielding a stationary, powerlaw–like distribution and under-threshold survival. Pg–Vp co-cultures further demonstrate replicate *rescue* outcomes for subcritical inocula. Critically, Vp reliably saturates to capacity, constraining terminal phases within the experimental horizon to *coexistence* (Pg persists with Vp at capacity) or *Pg extinction*. A two-species replicator model maps these outcomes onto a **(*β, γ*)** plane, restricting accessible regions once Vp is established and suggesting interventions that reduce facilitation or variability to restore eubiosis and limit Pg-associated inflammation.

## 1 Introduction

Periodontitis is among the most prevalent chronic inflammatory diseases, affecting nearly half of the global adult population [1]. Beyond local tissue destruction and tooth loss, periodontitis is now recognized as a systemic disease modifier that contributes to cardiovascular disorders, metabolic dysregulation, adverse pregnancy outcomes, and neurodegenerative conditions [2–4]. Despite this broad health burden, the microecological and biophysical principles that allow periodontal pathogens to persist and destabilize microbial communities remain incompletely understood [5, 6].

As a multifactorial, biofilm-driven condition, periodontitis emerges through a shift in the oral microbiota from a commensal-dominated, health-associated state (eubiosis) to a pathogen-enriched, pro-inflammatory configuration (dysbiosis) [6, 7]. These contrasting states can be understood as alternative stable equilibria of the microbial community. Eubiosis reflects a balanced composition that resists pathogen overgrowth and supports immune homeostasis, whereas dysbiosis is characterized by the enrichment of *Porphyromonas gingivalis* (Pg) and other pathogens, promoting sustained inflammation and tissue destruction [8].

Central to this transition is *P. gingivalis*, a Gram-negative obligate anaerobe that, even at low relative abundance, can remodel microbial community structure and modulate host immune surveillance [9, 10]. This observation, formalized in the keystone pathogen hypothesis, reframes pathogenicity in terms of ecological impact rather than absolute population dominance [11]. However, most studies have focused on individual virulence factors or host interactions in isolation, rather than examining how ecological thresholds, stochastic survival strategies, and cross-species interactions combine to determine *P. gingivalis* persistence [12–14].

A paradox emerges from this perspective, as *P. gingivalis* is frequently detected at low levels in both healthy and diseased conditions despite exhibiting Allee-type growth behavior [10, 15], a form of inverse density dependence in which population growth becomes negative below a critical threshold [16]. Classical population theory would therefore predict extinction of subthreshold populations, yet clinical studies report that *P. gingivalis* may persist, if at low abundance, in a fraction of individuals with periodontal health [17, 18] or reappear after antimicrobial or mechanical debridement [19–21]. This discrepancy implies that buffering mechanisms enable *P. gingivalis* to cross or bypass deterministic extinction thresholds and push the community toward the dysbiotic basin of attraction [14, 22].

One such mechanism may be microbial facilitation. In polymicrobial communities, metabolic interdependence and cooperative interactions are common [6, 14]. *Veillonella parvula* (Vp), an early colonizer of oral biofilms, co-occupies niches with *P. gingivalis* and may supply diffusible growth factors or alter redox conditions to favor *P. gingivalis* establishment [6, 15, 23]. *In vitro* studies support this hypothesis, showing that Vp-conditioned medium can significantly alter *P. gingivalis* growth dynamics and lower its effective colonization threshold [12, 15]. However, the magnitude and mechanism of this facilitation remain to be fully elucidated [23].

In addition to facilitation, stochastic effects stemming from microenvironmental variability may enable *P. gingivalis* populations to survive at densities where deterministic models predict extinction [9, 15, 24, 25]. Phenotypic heterogeneity, arising from gene-expression noise and micro-niche variation, has been shown to confer survival advantages through bet-hedging strategies [2, 14, 26]. For *P. gingivalis*, such heterogeneity could represent a crucial colonization tactic, helping push populations across the tipping point that separates extinction from dominance [26].

Lastly, the dynamics of colonization are shaped by nonlinear, frequency-dependent interspecies interactions. Evolutionary game theory offers a principled framework for describing how competition and cooperation generate alternative stable states in microbial communities [27, 28]. In the Pg–Vp system, cooperation may underlie dysbiotic transitions, whereas competition may correspond to eubiotic resilience where Pg fails to replicate [6].

In this study, we integrate quantitative experiments with mathematical modeling to address three questions: (i) How does *V. parvula* influence the colonization threshold of *P. gingivalis*? (ii) Can heterogeneity-driven noise promote *P. gingivalis* persistence below deterministic thresholds? (iii) Can we identify the Pg-Vp interactions that lead to experimentally observed dynamic phases (microbiome constellations)? To answer these, we parameterize an Allee-effect model, quantify Vp-driven threshold shifts, incorporate stochasticity to explore noise-induced rescue, and use a replicator equation, along with new Pg-Vp coculture experiments, to map potential bacterial interactions in their corresponding phases.

Together, these approaches establish a systems-level perspective on *P. gingivalis* microecology and provide a quantitative basis for explaining the persistence of this pathogen despite its strong Allee dependency. Understanding these ecological mechanisms is essential for designing targeted interventions that prevent *P. gingivalis* colonization or restore microbial homeostasis to mitigate the systemic consequences of chronic oral inflammation [6, 11].

## 2 Experimental Basis

Our modeling is based on quantitative growth data, focusing on three key experimental series reported by Hoare et al. [15], as well as new experiments designed to evaluate the effects of stochasticity on Pg dynamics. As reported by Hoare et al. [15], single-species inoculum experiments were performed by culturing Pg strain 381 across a wide range of starting densities (10^5^–10^8^ cells*/*mL). Cultures were incubated anaerobically at 37 °C in mucin–serum medium (2.5 mg/mL hog gastric mucin, 2.0 mg/mL proteose peptone, 2.5 mg/mL KCl, 1.0 mg/mL trypticase peptone, 1.0 mg/mL yeast extract, 1.0 µg/mL cysteine HCl, 10% heat-inactivated human AB serum, 5 µg/mL hemin). Growth was monitored daily by qPCR targeting the 16S rRNA gene. These data were used to parameterize a cubic Allee-effect model capturing density-dependent deterministic dynamics. The resulting parameters then served as the baseline for our stochastic analysis of *P. gingivalis* population behavior, enabling us to examine how microenvironmental noise perturbs deterministic growth trajectories.

Second, we incorporated results from spent-medium complementation assays, in which cell-free supernatant from stationary-phase Vp cultures was added to low-density Pg inocula [15]. The spent medium of Vp was obtained by growing the bacteria in mucin–serum medium under anaerobic conditions for 24 hours. The cultures were then centrifuged, and the resulting supernatants were filter-sterilized twice using 0.22 µm filters to ensure removal of all cells. This sterile, cell-free spent medium was subsequently used at different proportions mixed with fresh medium, followed by inoculation of Pg at low-cell-density (10^5^ cells*/*mL). While the experimental observations indicated rescue from extinction, model fitting allowed us to quantify the extent of threshold reduction through changes in the Allee parameter (*A*).

Third, Pg and Vp were co-inoculated into mucin–serum medium at a low cell density of 10^5^ cells/mL each, and cultured under anaerobic conditions in a batch setup [15]. The cultures were incubated at 37 °C and sampled daily. The growth of Pg was quantitatively assessed using qPCR. This experimental design aimed to evaluate interspecies interactions between early colonizers like Vp and Pg. The observed growth of Pg from this co-culture experiment was used to inform and parameterize the explanatory two-species replicator model, which captures the frequency-dependent interactions and fitness landscape between the two species.

Fourth, to experimentally assess model predictions related to the influence of stochastic fluctuations, we conducted a new set of long-term monoculture experiments in which Pg was inoculated at a low cell density of 10^5^ cells/mL in mucin–serum medium and monitored for up to 32 days (27 biological replicates). Cultures were sampled inside the anaerobic chamber, followed by serial dilutions and plating on

Brain Heart Infusion agar supplemented with blood. Colony-forming units (CFUs) were determined after 7 days of growth. These experiments also included co-cultures of Pg with Vp, with the latter either added at the beginning of the experiment as a co-culture control, or introduced after Pg had been incubated for 9 days to evaluate whether *P. gingivalis* populations could be rescued from extinction.

## 3 Critical *P. gingivalis* Density Threshold: Experimental Evidence for Allee Effect

A central question in oral microbial ecology is why *Porphyromonas gingivalis* (Pg) can persist at low levels in some individuals but becomes extinct in others. To address this, we modelled *P. gingivalis* population growth using a cubic Allee-effect differential equation, which encapsulates the principle that populations must surpass a critical quorum to sustain growth [29]. The model equation reads:

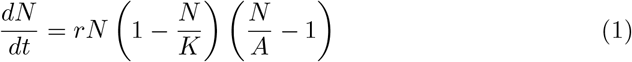

Here, *N* (*t*) is the *P. gingivalis* population (gene copies*/*ml) at time *t, r* is the intrinsic growth rate, *K* is the carrying capacity, and *A* is the Allee threshold. The last term encodes inverse density dependence, when *N < A* growth becomes negative and the population heads toward extinction.

This model was fitted to experimental time-series data [15] using the differential evolution algorithm [30].

Fitting this model to experimental growth curves yielded *r* = 0.6529, *K* = 2.08 × 10^9^ ± 2.02 × 10^5^, and *A* = 4.59 × 10^7^ ± 2.6 × 10^3^, closely reproducing experimental observations (Fig. 1. Please note that the Pg decline after reaching the theoretical capacity can be mediated by nutrient depletion or metabolic waste buildup [31], phenomena that are not captured by our rudimentary model. These results provide quantitative evidence that *P. gingivalis* requires a minimal quorum to escape extinction, consistent with microbial ecological theory.

**Fig. 1:**
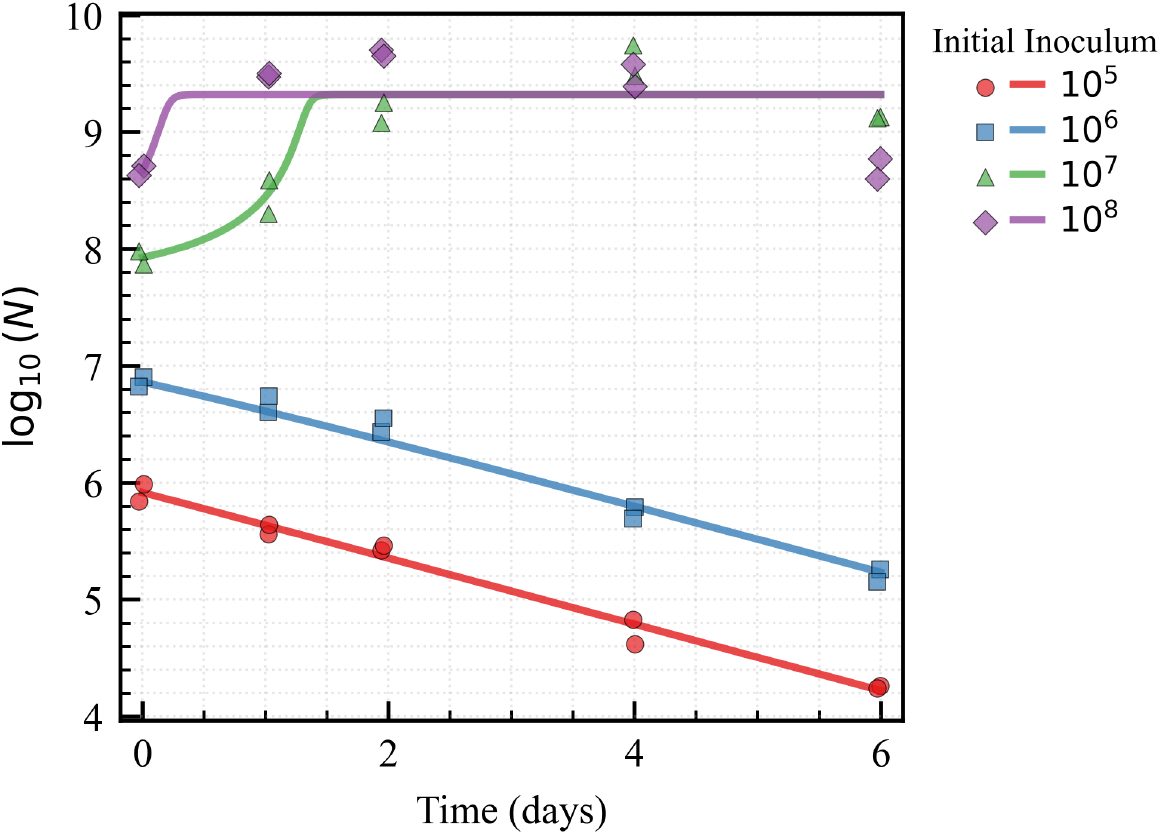
Model fitting results across four initial inoculum sizes. **Scatter points** represent observed values of log_10_(*N*) over time, while **solid lines** show predictions from the fitted model. Each color/marker denotes a distinct initial 10^x^ inoculum level.

This threshold shows why *P. gingivalis* is rarely sustained in healthy oral microbiomes. Host immune clearance and competitive commensals keep *P. gingivalis* below the *A* threshold. Once the threshold is crossed, however, *P. gingivalis* growth becomes self-reinforcing, establishing a dysbiotic community that is difficult to reverse.

## 4 *V. parvula* -Derived Metabolites Facilitate *P. gingivalis* Persistence at Subcritical Densities

Because the estimated Allee threshold *A* is high relative to *P. gingivalis* abundances in periodontal health and even moderate periodontitis, we asked to what extent facilitation by early colonizers lowers this barrier. We therefore quantified the effect of diffusible *V. parvula* (Vp) metabolites on the effective threshold by growing *P. gingivalis* in media supplemented with cell-free Vp-conditioned supernatant harvested after 24 h and re-estimating *A* under these conditions [15].

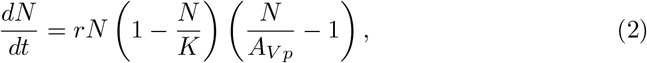

The Allee model was re-fitted to the Pg-Vp spent medium data where *A*_V p_ is the metabolite-modified threshold. The fitted results showed a significant reduction in *A*_V p_ compared to previously estimated *A* across all tested spent medium concentrations (Fig. 2b). This suggests that Vp-derived metabolites lower the minimum population size required for *P. gingivalis* persistence.

**Fig. 2:**
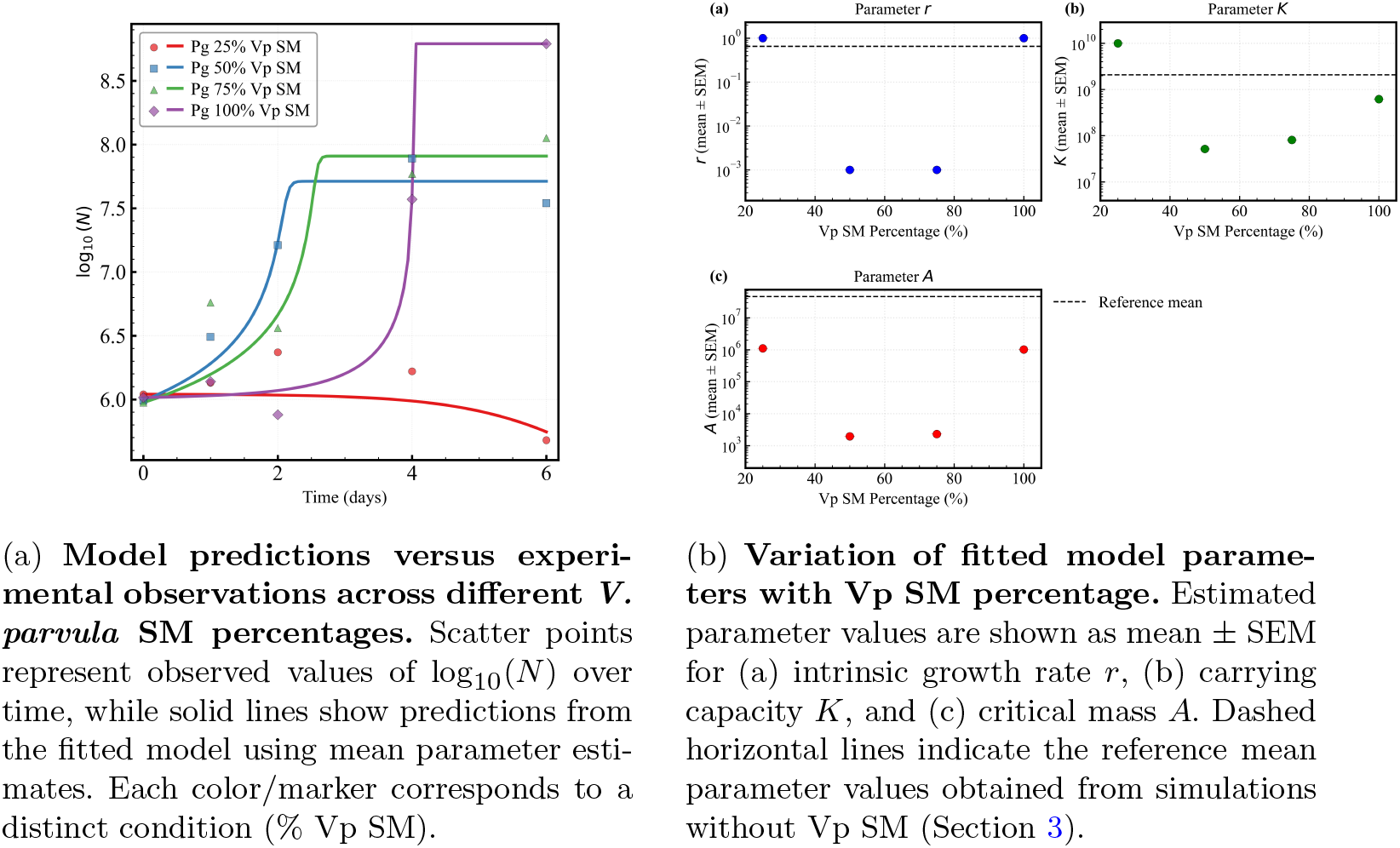
Effect of Vp-derived spent medium on *P. gingivalis* population growth and fitted parameters.

Vp-conditioned medium lowered the fitted Allee threshold relative to baseline (*A*_V p_ *< A*), but the effect was condition-dependent. For example, as shown in Fig. 2a, at 25% Vp SM, subcritical *P. gingivalis* inocula failed to grow, consistent with the fitted *A*_V p_ = 1.1 × 10^6^ ± 4.09 × 10^−1^ remaining above the starting density of 1.09 × 10^6^ (Figs. 2b-c, 2a). Across SM levels, estimates of *r* and *K* also varied (Fig. 2b), but the key factor for Pg growth persistence was the shift in *A*, which defines the extinction–persistence boundary. These parameter variabilities, together with the recurrent detection of low-abundance *P. gingivalis in vivo*, motivate a stochastic extension in which *A* and *r* stochastically fluctuate to model microenvironmental and demographic noise.

## 5 Microenvironmental Noise Enables *P. gingivalis* Persistence

The deterministic Allee model captures how *V. parvula*–derived metabolites lower the Pg persistence threshold, but it cannot explain replicate-to-replicate variability near this barrier. *In vivo*, however, *P. gingivalis* frequently persists at low abundance, suggesting that microenvironmental variability and cellular heterogeneity intermittently push populations across the deterministic threshold.

To capture this variability, we incorporated environmental and demographic noise into the model by allowing the threshold (*A*) and proliferation rate (*r*) to fluctuate. These stochastic terms collectively account for variability in microenvironmental conditions and cell-to-cell heterogeneity affecting proliferation and sensing behavior.

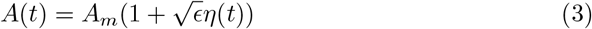

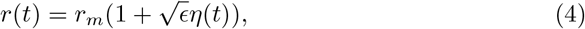

where *η*(*t*) is Gaussian white noise, with ⟨*η*(*t*)⟩ = 0 and ⟨*η*(*t*)*η*(*t*^*′*^)⟩ = *δ*(*t*−*t*^*′*^), and *ϵ* controls the overall noise intensity. *A*_m_ and *r*_m_ correspond to the baseline deterministic (mean) Allee threshold and proliferation rate, respectively. For analytical tractability, we have applied the same noise term in both parameters. Substituting into the Allee model yields:

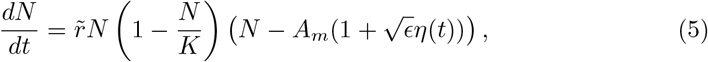

where 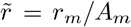 is a rescaled proliferation rate. We interpret the above equation in the Itô sense; *N* = 0 (extinction) and *N* = *K* (capacity) are natural/absorbing boundaries unless stated otherwise. This results in the stochastic differential equation:

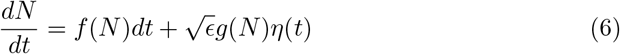

with:

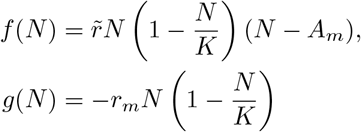

Assuming small noise intensity *ϵ* ≪ 1, we can apply Novikov’s theorem [32, 33] to obtain an extended mean-field approximation:

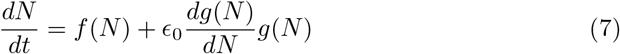

The term *ϵ*_0_ *g*^*′*^(*N*)*g*(*N*) represents the mean-field correction for multiplicative noise. Since we interpret Eq. (6) in the Itô sense, we treat *ϵ*_0_ as an effective fitted coefficient. After some algebra, the mean-field equation reads:

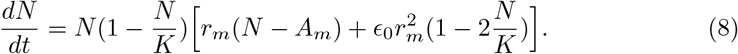

As one observes, the third term on the right-hand side allows us to calculate the new saddle node, which depends on the noise intensity:

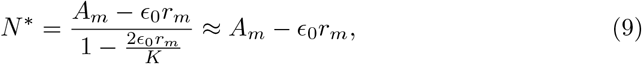

since the term 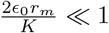. Therefore, noise shifts the saddle node, i.e., Allee threshold, towards zero (see Fig. 3). Thus, minimal quorum-sensing noise lowers the effective Allee barrier by ≈ *ϵ*_0_*r*_m_, increasing the chance of under-threshold rescue. This approximation is valid only for small noises, as a prerequisite for Novikov’s theorem.

**Fig. 3:**
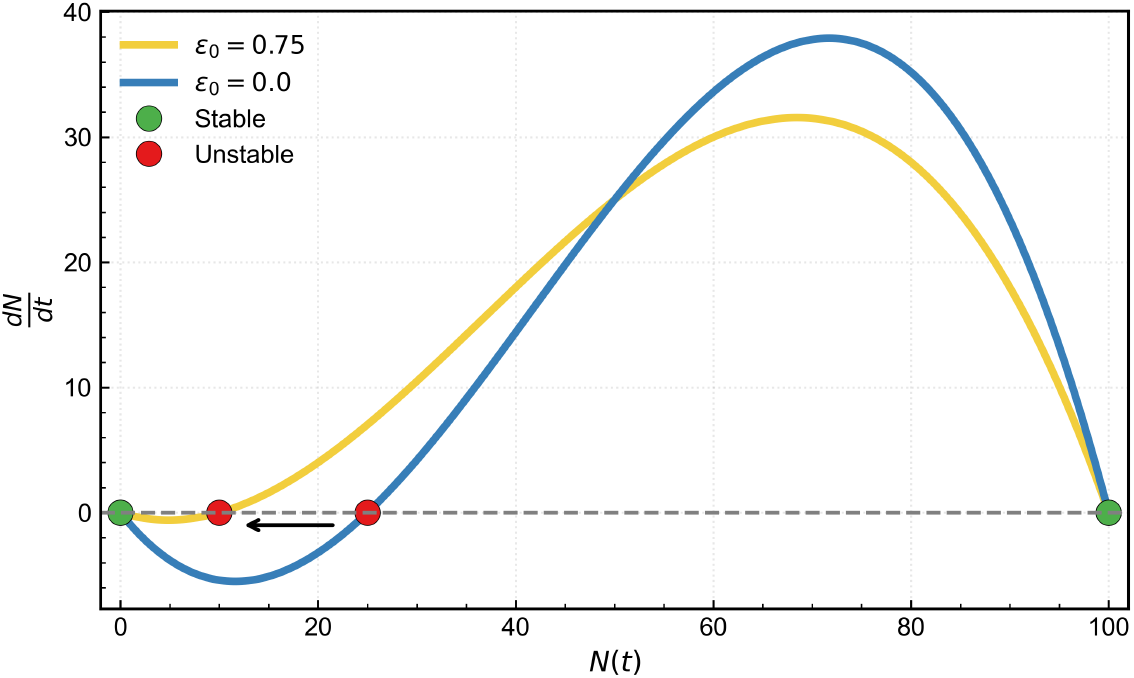
Illustrative nullcline plot of the bacterial population *N* (*t*) under two noise strengths. Parameters (*K* = 100, *A* = 25, *r* = 1) were chosen for demonstration purposes to clarify the qualitative effect of noise on the Allee threshold.

At higher noise amplitudes (*ϵ* ≫ 1), populations initiated above the Allee threshold (*N*_0_ *> A*) remained robust to stochastic fluctuations and maintained populations near carrying capacity (Fig. 4a). In contrast, when populations started below the threshold (*N*_0_ *< A*), stochastic variability occasionally enhanced survival by driving trajectories above the deterministic threshold but without reaching carrying capacity, or sustaining persistence at low abundances, thereby rescuing populations from extinction (Fig. 4b). To validate the predictions of the stochastic model, we first conducted long-term experimental observations of subthreshold *P. gingivalis* cultures. We then derived the corresponding steady-state probability distribution using the Fokker–Planck equation and compared it with the long-term subthreshold laboratory cultures of Pg.

**Fig. 4:**
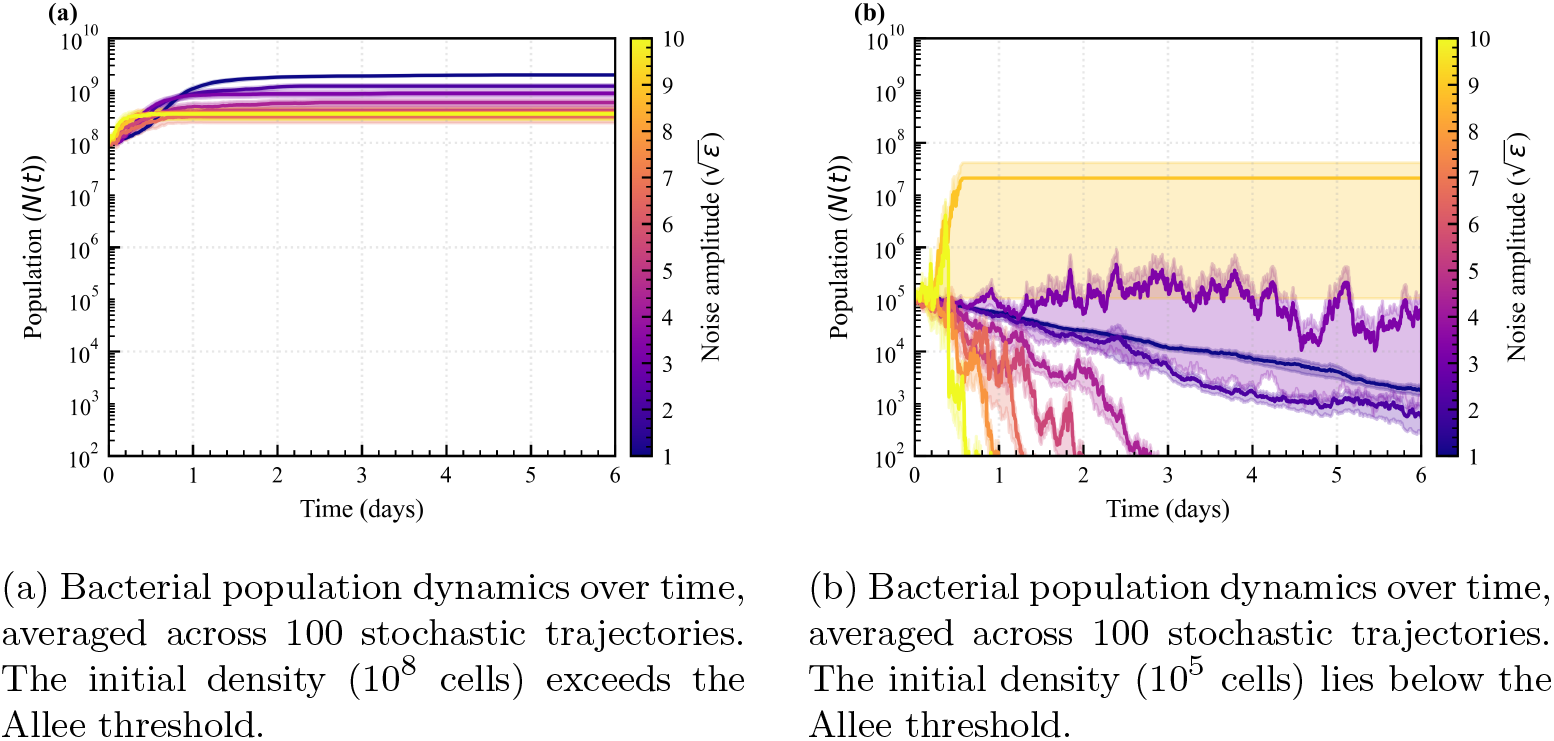
Stochastic simulations of bacterial population dynamics under different perturbation strengths. Each curve represents the mean across 100 independent stochastic trajectories, with shaded regions denoting the standard error of the mean. Model parameters were obtained by fitting an Allee-effect growth model to experimental growth curves (Section 3): carrying capacity *K* = 2.08 × 10^9^, Allee threshold *A* = 4.59 × 10^7^, and intrinsic growth rate *r* = 0.6529. The stochastic term was implemented as Gaussian white noise with mean 0 and standard deviation 1.

Figure 5 shows the experimentally measured *P. gingivalis* trajectories over time and the corresponding steady-state probability distribution *P*_eq_(*N*), based on observations up to 32 days across 27 biological replicates. The data reveal heterogeneous steady-state outcomes, with some populations collapsing to very low abundances (*N* ≲ 10^2^) at different rates, while others maintain higher population levels (*N* ≳ 10^5^) after fluctuating around the threshold.

**Fig. 5:**
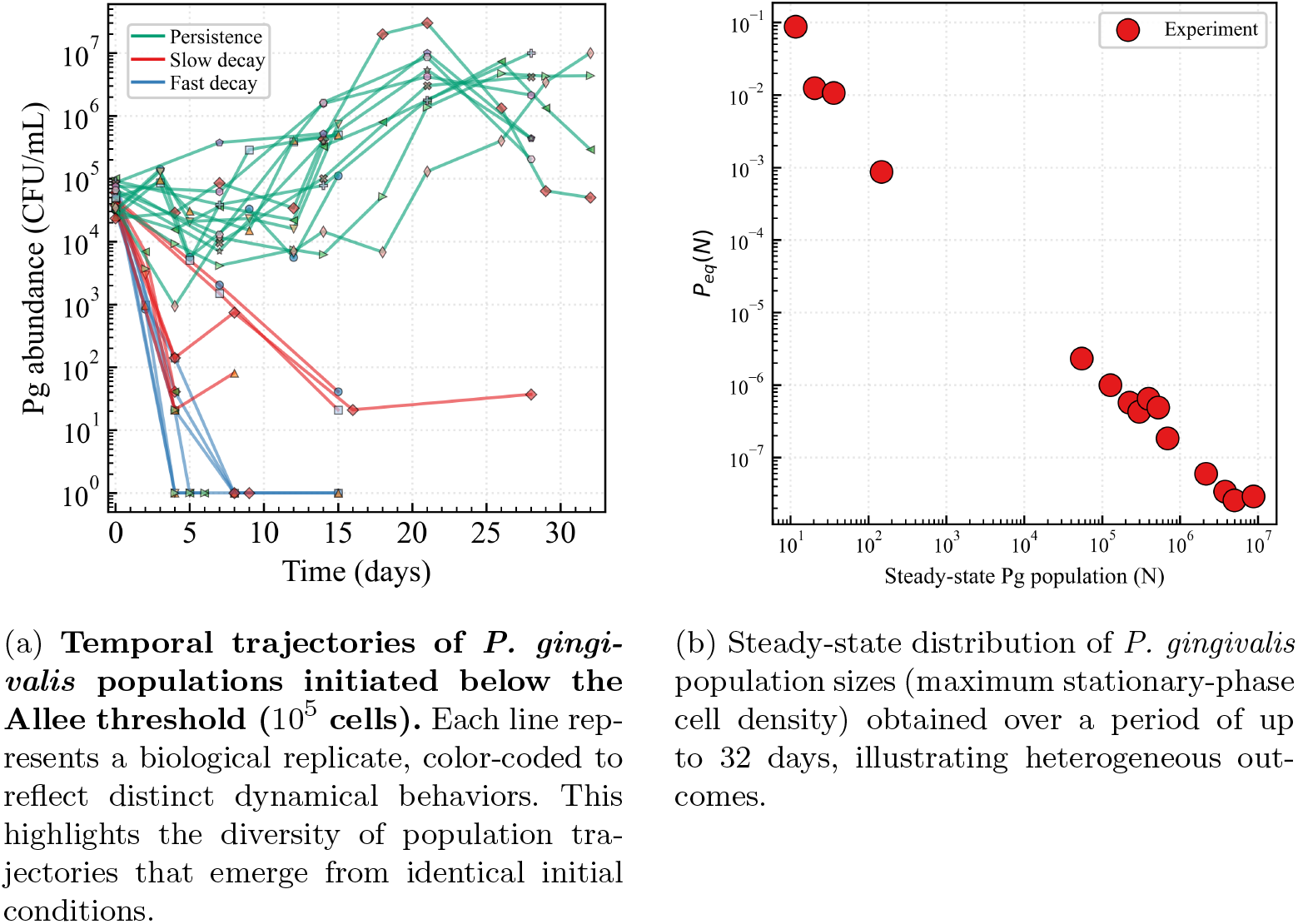
Experimental validation of the stochastic persistence model for *P. gingivalis* subthreshold populations. Both panels summarize data from 27 biological replicates: (a) the steady-state population size distribution observed within a 32-day period, and (b) the corresponding temporal trajectories showing stochastic transitions between extinction and persistence states.

This diversity of outcomes is consistent with the stochastic model predictions under intermediate noise intensities (3 ≤ *ϵ* ≤ 5), where random fluctuations can drive populations either toward extinction or persistence. The limited number of observations at intermediate abundances (10^2^ *< N <* 10^5^) suggests that threshold-crossing events occur over short timescales, with populations spending relatively little time in this transition region. As shown in Fig. 5a, individual trajectories exhibit such noise-driven fluctuations near the Allee threshold before stabilizing at low or high abundance levels. Together, these results demonstrate strong agreement between the experimental observations and the stochastic model, supporting the interpretation that *P. gingivalis* population dynamics are governed by stochastic transitions between extinction and persistence within a metastable landscape.

For a quantitative link between stochastic dynamics and population persistence, we computed the temporal evolution and steady-state probability distribution of the *P. gingivalis* population from the corresponding Fokker–Planck equation (FPE) derived from Eq. (6).

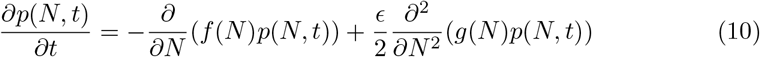

The equilibrium distribution for the multiplicative noise FPE reads:

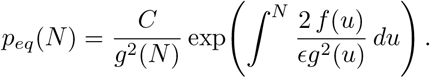

The resulting stationary probability density *p*_eq_(*N*) captures the main features of the experimentally measured frequency distribution obtained from 27 independent replicates initiated below the deterministic Allee threshold.

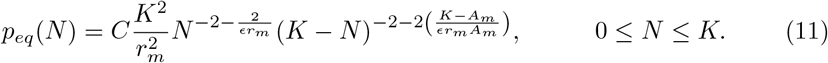

The steady-state distribution *p*_eq_(*N*) resembles a Beta-like distribution with compact support between [0, *K*], consistent with the bounded nature of bacterial population sizes. For our Pg growth experiments, a truncated power law trend *p*_eq_(*N*) ∝ *N*^−α^ was observed near the extinction state (*N <* 10^2^) with an estimated exponent *α* = 2.73 ± 0.46. Plugging into our theoretical exponent 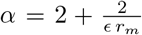 the experimentally obtained value, we find that the *ϵ* ≥ 2.6. Unfortunately, fitting the full distribution is difficult since we do not have observations close to the capacity. The fitted *ϵ* reflects the minimal intrinsic noise level that enables under-threshold populations to persist (see Fig. 5b). Biologically, the inferred *ε* quantifies a minimal microenvironmental-sensing noise sufficient to rescue underthreshold inocula; *in vivo*, additional host and community variability will further increase effective noise, raising the probability of escapes across the Allee barrier.

In summary, incorporating noise into the Allee framework reveals that random fluctuations can shift the effective persistence threshold and rescue populations that would otherwise collapse. This stochastic mechanism provides a quantitative explanation for the intermittent detection of *P. gingivalis* at low abundance *in vivo* and indicates that persistence is not purely deterministic. Instead, phenotypic heterogeneity and micro-niche variability likely act as bet-hedging strategies that promote long-term survival.

## 6 Microecological Phase Transitions Driven by Pg–Vp Interactions

To investigate how pairwise interactions shape microecological dynamics, we focused on a system comprising *P. gingivalis* (Pg) and *V. parvula* (Vp). This approach allows us to explore fundamental ecological behaviors that may underlie more complex community transitions.

For mechanistic modeling purposes, we describe the possible outcomes of the Pg– Vp system using four terms: **Pg extinction, Pg dominance, stable coexistence**, and **bistability**. These terms capture the distinct dynamical states predicted by our model and reflect alternative ecological attractors that can emerge from pairwise interactions.

Let *N*_Pg_ and *N*_Vp_ denote the absolute abundances of *P. gingivalis* and *V. parvula*, respectively, and *N*_total_ = *N*_Pg_ + *N*_Vp_ the total number of bacteria. We define the normalized relative abundance of Pg as:

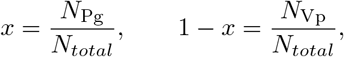

so that *x* ∈ [0, 1] fully characterizes the community composition. The four qualitatively distinct outcomes can then be described as follows:

- **Pg extinction:** Pg is either absent or remains at low relative frequency, and the community resists Pg invasion (*x* = 0 is a stable equilibrium).
- **Pg dominance:** Pg proliferates and becomes dominant, leading to a Pg-dominated community (*x* = 1 or a Pg-dominated interior equilibrium is stable).
- **Stable coexistence:** An interior equilibrium 0 *< x*^⋆^ *<* 1 exists and is stable, indicating that Pg and Vp persist together at steady state.
- **Bistability:** Both *x* = 0 (Pg extinction) and *x* = 1 (Pg dominance) are stable equilibria, separated by an unstable interior threshold *x*^⋆^. The final outcome depends on initial conditions, reflecting historical contingency in community assembly.

Each species’ fitness is determined by its interactions with Pg and Vp, encoded in the payoff matrix

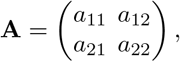

where *a*_ij_ is the payoff to species *i* when interacting with *j*. The expected fitnesses are

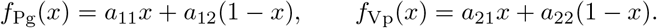

The replicator equation is the dynamical core of evolutionary game theory that translates payoff matrices into population dynamics [34, 35]. Here, the replicator equation states that the rate of change of *x* is proportional to Pg’s excess fitness over the population mean:

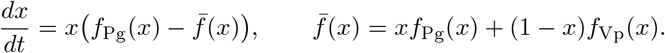

For two species, this simplifies to

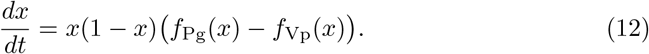

**Table 1:** Biological interpretation of the replicator dynamics across the (*β, γ*) plane for *x* ∈ [0, 1] (cf. Fig. 6).

| Region | Condition | Attractors<sup>a</sup> |
| --- | --- | --- |
| Pg extinction | $\beta < 0, \beta + \gamma < 0,$ | $x = 0$ stable |
| Pg dominance | $\beta > 0, \beta + \gamma > 0,$ | $x = 1$ stable |
| Coexistence | $\beta > 0, \beta + \gamma < 0$ | $x = 0$ and $x = 1$ unstable; $x^*$ stable |
| Bistable | $\beta < 0, \beta + \gamma > 0$ | $x = 0$ and $x = 1$ stable; $x^*$ unstable |

<sup>a</sup> $x^*$ denotes the interior fixed point (when it exists).

Substituting the linear fitness functions gives:

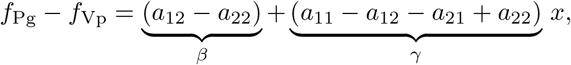

and hence the normalized Eq. (12) reduces to a single cubic ODE:

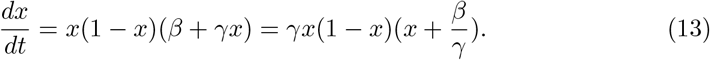

The parameter *β* quantifies Pg’s relative advantage when interacting with Vp (against Vp with itself) and *γ* compares self-interaction against cross-species fitness changes. The system has three equilibria: (*x* = 0) (Pg extinction), (*x* = 1) (Pg dominance), and (*x*^⋆^ = ™ *β/γ*) (an interior tipping point if (0 *< x*^⋆^ *<* 1)). Stability analysis shows that (*x* = 0) is stable when (*β <* 0), (*x* = 1) is stable when (*β* + *γ >* 0), and (*x*^⋆^) is an unstable saddle if (*γ >* 0). On the other hand, the (*x*^⋆^) can become stable when *β <* 0 and *β* + *γ >* 0. Table (1) resumes the biological interpretation of our stability analysis results. Using the stability conditions of *γ* and *β* parameters for the replicator Eq. (13), one can compute a phase space diagram as shown in Fig. 6 where four phases/attractors lie (Extinction, Dominance, Coexistence, and Bistability state), separated by two boundaries (i.e., *β* + *γ* = 0 and *β* = 0).

**Fig. 6:**
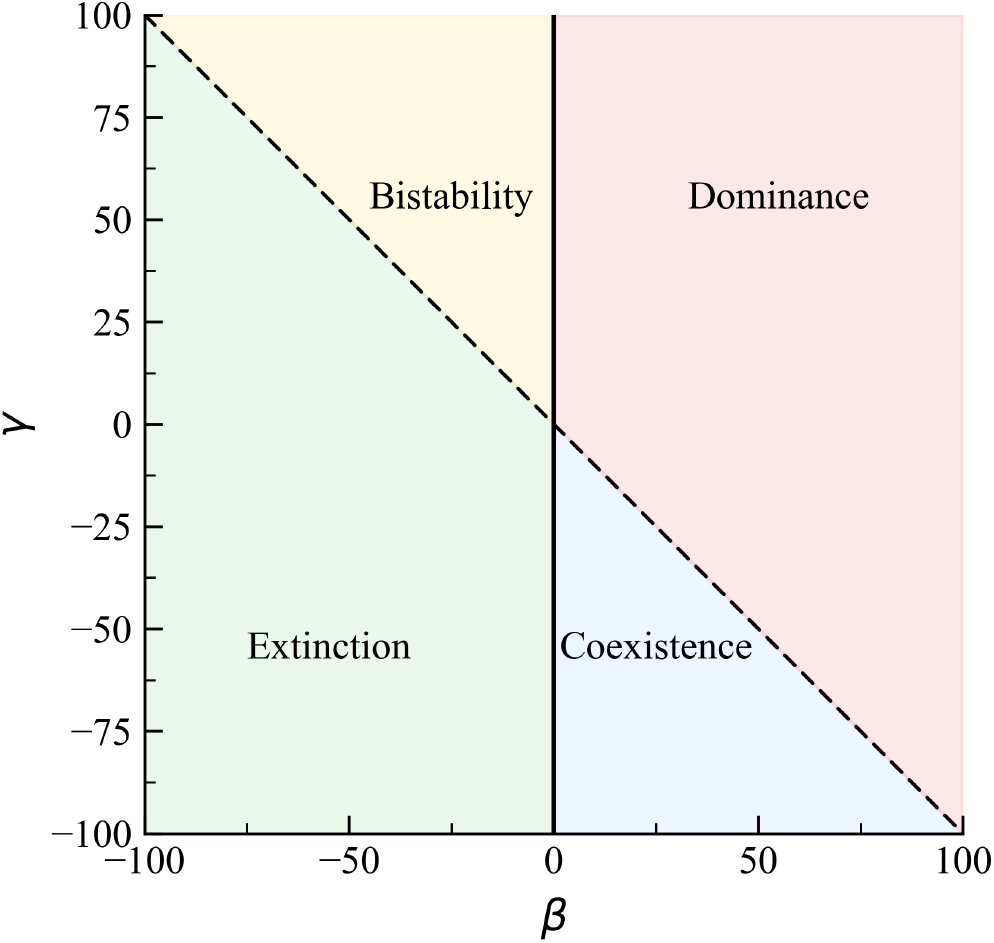
Phase space plot of Pg-Vp interactions. Here *β* + *γ* = 0 is shown in dotted line and *β* = 0 is shown in solid line.

To constrain our theoretical results, we performed Pg-Vp co-culture experiments under a subcritical inoculum (*N* (0) = 10^5^). When *V. parvula* was co-cultured with *P. gingivalis* on Day 0, all replicates exhibited sustained growth, reaching near carrying capacity levels, consistent with strong early cross-species facilitation (Fig. 7a). In contrast, when *V. parvula* was introduced only after the onset of population decline (Day 9), the outcomes varied. In several replicates, Pg regained growth to high densities following several days of sustained population levels (‘Persistence’), whereas in others, the population was stochastically rescued (“Rescue”) or sometimes went extinct despite Vp addition. Notably, when *P. gingivalis* was cultured alone before adding Vp, decline occurred in all replicates but at variable rates, consistent with the stochastic model predictions of noise-driven variability in time to extinction (Fig. 4b).

**Fig. 7:**
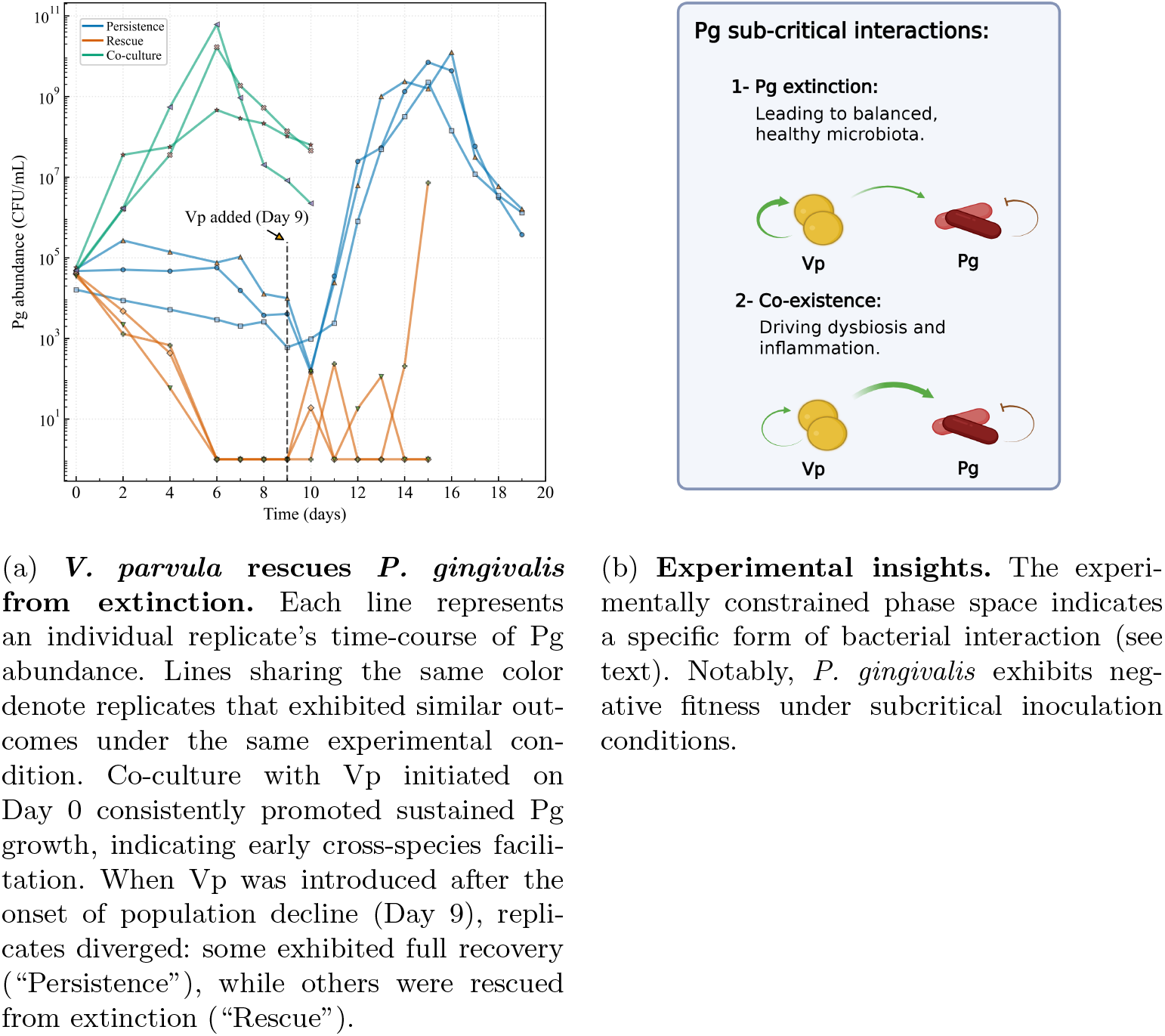
Our Pg–Vp co-cultures demonstrated the expected facilitation of Pg growth. These experimental results constrain the phase space of Pg–Vp outcomes and imply experimentally testable interactions between Vp and Pg.

By the end of all co-culture assays, Vp consistently reaches (or closely approaches) its carrying capacity (own experimental data). Although Vp time series are not available, imposing restrictions on the identifiability of the payoff matrix, this terminal saturation imposes a strong phase constraint: Pg-dominant endpoints are not observed. Within the experimental horizon, attainable terminal states reduce to two possibilities: (i) *coexistence*, with Pg at positive abundance while Vp occupies (near-)capacity; or (ii) *Pg extinction*, with Vp alone persisting. Stochasticity blurs the separatrix between these outcomes, producing the observed replicate variability. This experimental constraint on our phase space events has interesting implications for Pg-Vp interaction payoff parameters. Our experiments suggest *β* + *γ <* 0 =⇒ *α*_11_ *< α*_21_. Assuming that subcritical inoculation of Pg implies *α*_11_ *<* 0, no Pg-on-Vp interaction needs to be assumed. If the Vp facilitation on Pg is larger than the contribution of Vp selfrenewal, i.e. *β >* 0 =⇒ *α*_12_ *> α*_22_, it leads to bacterial coexistence. The latter can explain the Pg rescue in the presence of Vp (even when added at later times), since the Pg elimination state *x* = 0 becomes unstable. Fig. 7b schematically recapitulates our findings.

## 7 Discussion

Dysbiosis in periodontitis is increasingly recognized as a microecological phase transition in the oral microbiome, in which a polymicrobial community enriched with pathobionts such as Porphyromonas gingivalis emerges and disrupts the balanced commensal community [9, 36]. Our results unify experimental observations and theory by demonstrating that *P. gingivalis* persistence emerges from the interplay of (i) density-dependent growth with an Allee threshold, (ii) cross-species facilitation by *V. parvula*, and (iii) stochastic fluctuations possibly induced by environmental and phenotypic heterogeneity. Together, these mechanisms explain how *P. gingivalis* can persist at low densities *in vivo* despite deterministic predictions of extinction (see Fig. 8).

**Fig. 8:**
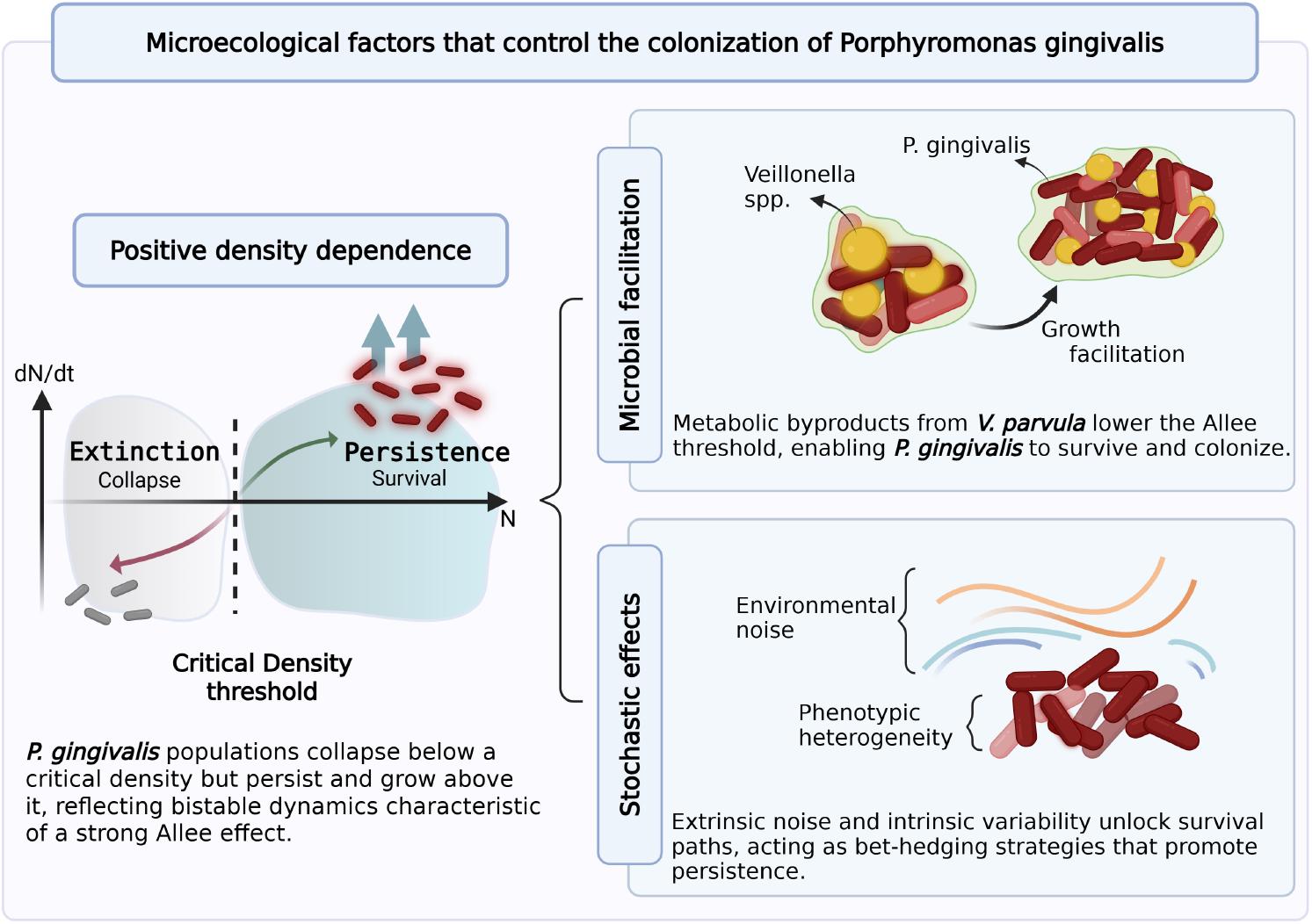
Overview of ecological processes influencing *P. gingivalis* persistence. Population survival depends on crossing a minimum density, below which extinction occurs and above which growth is maintained. Early colonizing species, such as *V. parvula*, provide metabolic support that allows *P. gingivalis* to establish even when present at low abundance. Random fluctuations in the environment, as well as intrinsic differences among individual cells, create variability that increases the likelihood of survival under adverse conditions.

Our deterministic Allee model quantitatively confirms the existence of a critical density threshold (*A*) below which *P. gingivalis* populations collapse, consistent with microenvironmental-sensing theory and the cooperative traits required for virulence and biofilm maturation [16, 37]. Importantly, the addition of Vp-conditioned medium lowered this threshold, supporting the idea that *V. parvula* functions as an ecological enabler that metabolically primes the niche for *P. gingivalis* colonization [15, 24].

While the metabolic cues required for Pg to support its own growth or those provided by Vp to Pg have not been identified yet, our work supports earlier findings which suggested Veillonella has a central role in the establishment of multi-species dental plaque communities [25, 31, 38]. Veillonella are prevalent dental plaque colonizers, previously recognized as a “core” subgingival species, which are proposed as generalists that play a central metabolic role supporting eubiosis to dysbiosis microbiome shifts Incorporating stochasticity revealed that noise can rescue subcritical populations by driving them across the unstable manifold toward persistence [39–41]. This result reframes *P. gingivalis* detection at low levels as a natural outcome of noise-driven survival and bet-hedging strategies [14, 42]. Additionally, by analyzing the corresponding FPE equilibrium distribution, we could obtain a quantitative noise estimate that is in line with the observed frequencies of sub-threshold survival. Clinical and experimental evidence supports this view: Few *P. gingivalis* cells can survive intracellularly [43, 44] and form persister subpopulations that tolerate antibiotics and re-initiate growth when stress abates [45–47]. Although our model does not explicitly resolve single-cell behavior, we hypothesize that fluctuations in cue thresholds, signal concentrations, or cellular responsiveness could generate phenotypic variability. Such heterogeneity may increase the likelihood of persistence at low densities, linking molecular variation to population outcomes.

Finally, our exploratory game-theoretic model captures the context-dependent switch between competitive and cooperative regimes. Mapping Pg–Vp dynamics in the (*β, γ*)-plane shows that Pg extinction corresponds to a globally attracting *x* = 0 state when *β <* 0, whereas Pg replication dominates when *β* + *γ >* 0. The bistable regime highlights the potential for history-dependent outcomes: once the community is perturbed across the tipping point *x*^⋆^ = ™ *β/γ*, it will commit to Pg-dominance, even if the parameters remain unchanged. This provides a quantitative framework for understanding why Pg persistence can be difficult to reverse once established. In coculture, subcritical Pg inocula exhibited replicate-to-replicate variable outcomes, as expected for noise-assisted barrier crossing in a facilitated background. Notably, Vp reliably saturated to (near) carrying capacity by the end of the experiment; as a result, Pg-dominant endpoints were not observed, and the attainable terminal phases within the experimental horizon were reduced to two: *coexistence* (Pg persists with Vp at capacity) or *Pg extinction*. The coexistence phase can be interpreted as a proxy for the oral dysbiosis state. In our experiments, we mainly observe Pg-Vp coexistence, since even when Pg is at very low abundance, Vp introduction can still destabilize the zero Pg state.

Translationally, these findings suggest that therapeutic strategies should focus on shifting *β* to negative values, thereby restoring eubiosis as the global attractor. Potential approaches include (i) promoting competitive colonizers to reduce *P. gingivalis* fitness (*a*_12_), (ii) disrupting metabolic facilitation pathways provided by *V. parvula*, (iii) enhancing host immune clearance of *P. gingivalis*, and (iv) timed ecological reseeding following biofilm disruption. These strategies can be systematically tested by monitoring longitudinal *P. gingivalis* trajectories and inferring effective (*β, γ*) values before and after the intervention.

The Fokker–Planck equation analysis provides a probabilistic view of *P. gingivalis* persistence, revealing that stochastic microenvironmental-sensing noise can sustain subcritical populations that would otherwise collapse deterministically. This noise is expected to be enhanced in *in vivo* settings, where microenvironmental complexity is much higher than in *in vitro* experiments. This observation bridges ecological modeling with the physics of decision-making in biological collectives: the bacterial population continuously samples its microenvironment, and random fluctuations in signaling effectively explore multiple “choices” between extinction and growth. Such stochastic switching mirrors cell-fate selection in multicellular systems, where intrinsic noise drives transitions between phenotypic attractors.

The cell decision-making aspect allows us to invoke the Least Microenvironmental Uncertainty Principle (LEUP) [48–52], where multicellular communities tend to minimize uncertainty about their microenvironment by adopting states that reduce phenotypic entropy. Within this framework, the switch between Pg extinction and Pg dominance can be viewed not only as a compositional phase transition but also as a collective decision-making process: facilitation by *V. parvula* reduces environmental uncertainty, stabilizing a Pg-permissive niche, whereas interventions that enhance competition increase uncertainty for Pg and thereby promote extinction. This interpretation situates our results within a broader decision-theoretic framework that links microbial ecology with general principles of cellular adaptation.

Our study entails some limitations. First, the game-theoretic model is parameterized using *P. gingivalis* trajectories alone, which underdetermines the full payoff matrix. Direct co-culture experiments measuring both *P. gingivalis* and *V. parvula* abundances are needed to fully resolve the bacteria interactions. Second, we used a linear payoff structure, which cannot capture saturating metabolic interactions or context-dependent facilitation. In particular, the observed Pg decline after reaching the theoretical capacity indicates nutrient depletion, metabolic imbalance, or any other Pg stressor [31] production that could be modelled by additional non-linearities. Third, our stochastic analysis employed a Gaussian noise approximation and small-noise closure, which may not accurately represent bursty or correlated fluctuations *in vivo*.

Finally, our model focuses on a two-species subsystem and ignores higher-order interactions with other oral taxa and host immune responses that could shift the location of the tipping point.

Future work should focus on quantifying the molecular basis of *V. parvula* facilitation, validating noise-induced persistence in controlled microfluidic systems, and calibrating the game-theoretic model with high-resolution longitudinal multi-omics data. Integrating these approaches will enable predictive control of Pg-associated dysbiosis and inform the rational design of microbiome-targeted therapies aimed at restoring oral and systemic health [6, 7].

## Computational Tools

All simulations and optimizations were conducted in Python 3.10. Core libraries included `NumPy` and `Pandas` for numerical operations, `SciPy` for integration and optimization, and `Matplotlib` for visualization. Custom implementations of the Euler-Maruyama scheme were used for stochastic integration. All code was modular, version-controlled, and is available upon request.

## Declarations

### Ethics approval and consent to participate

Not applicable.

### Consent for publication

All authors have read and approved the final manuscript for submission.

### Data availability

The datasets analyzed and/or generated during the current study are available from the corresponding author upon request.

### Materials availability

Not applicable.

### Code availability

All computational analyses and modeling code supporting this study are publicly available on GitHub at: https://github.com/Moemenhussein11/P. gingivalis-Colonization.

### Author contribution

Conceptualization: HH; Methodology: MH, AB, and HH; Experiments: MS, PID Investigation: MH and AB; Formal analysis: AB, MH; Writing—review and editing: MH, AB, MQ, MS, PID, and HH; Supervision: HH; Funding acquisition: HH, MQ, PID.

